# miEAA 2.0: Integrating multi-species microRNA enrichment analysis and workflow management systems

**DOI:** 10.1101/2020.03.05.978890

**Authors:** Fabian Kern, Tobias Fehlmann, Jeffrey Solomon, Louisa Schwed, Christina Backes, Eckart Meese, Andreas Keller

## Abstract

Gene set enrichment analysis has become one of the most frequently used applications in molecular biology research. Originally developed for gene sets, the same statistical principles are now available for all omics types. In 2016, we published the miRNA enrichment analysis and annotation tool (miEAA) for human precursor and mature miRNAs.

Here, we present miEAA 2.0, supporting miRNA input from *Homo sapiens, Mus musculus*, and *Rattus norvegicus*. To facilitate inclusion of miEAA in workflow systems, we implemented an Application Programming Interface (API). Users can perform miRNA set enrichment analysis using either the web-interface, a dedicated Python package, or custom remote clients. Moreover, the number of category sets was raised by an order of magnitude. We implemented novel categories like annotation confidence level or localisation in biological compartments. In combination with the miR-Base miRNA-version and miRNA-to-precursor converters, miEAA supports research settings where older releases of miRBase are in use. The web server also offers novel comprehensive visualisations such as heatmaps and running sum curves with background distributions. Lastly, additional methods to correct for multiple hypothesis testing were implemented. We demonstrate the new features using case studies for human kidney cancer and mouse samples. The tool is freely accessible at: https://www.ccb.uni-saarland.de/mieaa2.

## Introduction

Transcriptomics designates an indispensable set of techniques to study gene expression, often in a genome-wide manner, as the backbone of modern molecular biology and clinical research. The innumerable amount of classical bulk-sequencing datasets is further augmented by the recent advancements in high-resolution single-cell approaches. Since gene expression is constituted by many biological factors, experimental focus has been enlarged to include the regulatory non-coding transcriptome (ncRNAs), i.e. to RNA classes that regulate messenger RNAs (mRNAs) either directly or indirectly. Among these, microRNAs (miRNAs) are small non-coding RNAs, typically 18-25 nucleotides in length, loaded by proteins of the AGO-family to build RNA-induced silencing complexes (RISC) [1]. Gene regulation through the RISC complex is facilitated by one or two mature (−5*p*; −3*p*) miRNA arms, arising from one or several transcribed precursors [2]. Besides other modes of action, activated complexes target preferentially 3′-untranslated regions of mRNAs to induce either catalytic cleavage or translation repression. Hence, profiling miRNA expression contributes to the understanding of gene regulation and potentially portrays cellular states. To date, numerous studies highlight their informative role in disease detection, sub-type classification, or progression, such as for cancer [3], neurodegenerative [4], or metabolic disorders [5] with a variety of bio-specimens [6].

Considering that several thousands of miRNAs have already been discovered, many novel miRNA candidates have been additionally proposed [7], while the total number of human miRNAs is estimated to be 2, 300 [8]. Finding differences in expression for miRNAs is similar to mRNAs and therefore non-trivial. Differential gene expression studies often lead to dozens, hundreds, or even thousands of de-regulated genes. Thus, large scale studies often make use of the functionality of gene set enrichment analysis (GSEA) [9]. GSEA can further reduce large amounts of information towards a significant set of molecular functions, biological properties, or pathways of genes. In principle, a user inputs either a set or ordered list of genes and the tool runs the required statistical algorithms and provides background datasets to compare against.

Similar functionality was also implemented for other omics types, including proteomics, metagenomics, or epigenomics. An in-depth review of gene set analysis methods for data other than mRNAs demonstrates the increasing interest and demand of the community in respective tools [10]. We previously developed an approach tailored for both miRNA precursor enrichment and mature miRNA enrichment analyses, the miRNA enrichment analysis and annotation tool (miEAA) [11]. Here, we present an update of this tool that includes more categories, supports more organisms, has new statistical functionality and offers a standardised Application Programming Interface (API) to facilitate the inclusion of miEEA in modern data analysis workflows [12].

Given the growing interest in miRNAs, other tools with similar functionality to miEAA exist. Among the most functional tools, the recent successor version of TAM [13] introduced 1, 238 human miRNA set categories obtained from manual literature review of approximately 9, 000 scientific manuscripts, along with new query and visualisation features. In addition to the over- and under-representation analysis, users can compare the correlation of two miRNA lists under different disease conditions. All kinds of enrichment tools rely on high quality sets of miRNA categories that were either obtained by curation of scientific literature or collected from specific databases. For instance, curated miRNA annotations can be obtained from miRBase or miRCarta [14], miRNA-target interactions from miRTarBase [15], miRNA-pathway associations from miRPathDB [16], tissuespecific miRNAs from the human TissueAtlas [17], or miRNA-disease associations from HMDD [18] or MNDR [19], many of which were updated in the last two years. Further specialised annotations like miRNA and transcription factor interactions provided by TransmiR [20], miRNA sub-cellular localisations collected in RNALocate [21], or extra-cellular circulating miRNAs contained in miRandola [22] provide target categories for integrated enrichment analysis.

## MATERIALS AND METHODS

In miEAA 2.0 we provide support for 3 species (2 new), 24 new category sets, and updates to our pre-existing datasets. To unify data preprocessing, we implemented an automated pipeline using Snakemake [23], Python 3.6, and the pandas [24] Python package facilitating data collection and filtering steps. For each species and their corresponding data sources our pipeline performs the same basic process, consisting of downloading the datasets, cleaning and updating the miRNA and precursor identifiers, transforming the results into a Gene Matrix Transposed (GMT) file, and creating background reference sets. Files were copied to the web server without further modification.

### Data collection

Novel datasets were obtained to build our enrichment categories, consisting of Gene Ontology [25], miRTarBase 8.0 [15], KEGG [26], miRandola 2017 [22], miRPathDB 2.0 [16], TissueAtlas [17], MNDR v2.0 [19], NPInter 4.0 [27], RNALocate v2.0 [21], TAM 2.0 [13], and TransmiR v2.0 [20]. Other pre-existing datasets have been updated, including HMDD v3.0 [18] and miRBase v22.1 [28]. We retained the rest of our pre-existing datasets, namely miRWalk2.0 [29], our published age and gender dependent miRNAs and our distribution of miRNAs in immune cells [11]. All datasets contain miRNAs or precursors for *Homo sapiens*. When available, we also utilise the data for *Mus musculus* and *Rattus norvegicus*, allowing enrichment analysis on 39, 31, and 26 miRNA/precursor category sets, respectively. Raw datasets were obtained either through a direct download or via an API. In particular, the QuickGO and KEGG datasets are compiled by querying their corresponding REST APIs.

### Category data preprocessing

First, data from QuickGO was mapped back to miRBase using RNAcentral [30]. NCBI Gene was used in conjunction with miRTarBase to produce the indirect annotations. With the aid of the miRBaseConverter R package [31], miRNA and precursor names were translated to the latest version of miRBase. For KEGG Pathways and GO Annotations (direct and indirect through target genes from miRTarBase) we only keep miRNAs for which functional MTI support is available. In the MNDR diseases category set, we exclude HMDD data as it is precursor based, and MNDR is for mature miRNAs.

### Web server, statistics, and API implementation

The miEAA web server was built using a dockerized Django Web Framework v2.1, which exposes a web-API using the Django REST framework. The celery software was used as the job scheduler. Frontend libraries comprise Highcharts, dataTables, jquery, and Bootstrap. P-value correction methods were implemented using the R stats package. For the static GSEA running sum plots, a simulated background distribution is computed by randomly permuting the test set 100 times and traversing the running sum for each random permutation. Alongside our new API we provide a lightweight Python package, as well as a command line interface (CLI) tool, supporting Python 3.5 or higher. These are made freely available through the Python Package Index (pip) and through the *ccb-sb* conda channel.

### Case studies

Raw and reads per million miRNA mapping (rpmmm) normalized miRBase v21 precursor counts and metadata of kidney renal clear cell carcinoma case and control samples were obtained from TCGA. Since multiple sequencing results might be associated with the same sample ID in TCGA, we kept only one result file for each sample by preferring files from H over R over T analytes and selecting the aliquot with the highest plate number and / or lexicographical sorting order. Subsequently, miRNAs with fewer than 5 raw reads in less than 50% of either case or control samples were discarded from the analysis. All remaining miRNA counts were *log*_2_-scaled. Effect size was calculated using the implementation of Cohen’s d from the R package effsize. Lists of precursor names, either selected by statistical significance or ordered by effect size, were converted from miRBase v21 to v22.1 using the online miRBase converter feature of miEAA. The list of all precursors from miRBase v21, converted to v22.1, were used as a reference set. The configured parameters included default precursor category sets without the *PubMed ID* and *TransMiR Tissues* sets, BH-FDR adjustment to a significance level of 0.05 with independently adjusted p-values per category set, and a minimum of 2 required hits per sub-category.

For the second case study, raw Agilent microarray data along with sample metadata was downloaded from NCBI’s GEO using accession ID GSE117000. Array parsing and probe signal processing was performed identically to the description in the first publication of miEAA [11]. Subsequently, all counts were quantile-normalized and *log*_2_-transformed. All further down-stream analyses were performed analogous to the first case-study described above.

## RESULTS

### Overview on miEAA 2.0

In the following, changes and novelties introduced by the second major release of miEAA are described. Since all annotations of miRNAs to categories and databases are with respect to the miRNA reference database, miRBase, we converted the datasets to match its latest public version 22.1. This also affects the miRBase-version and miRNA-to-precursor converters, the former of which was designed to be fully backwards compatible. Moreover, both ORA and GSEA algorithms accept lists of either precursors or miRNAs, from human, mouse, and rat species. In total, 123, 655 categories from 15 published databases/resources are available to test against. A detailed breakdown of the counts by source and organism, on database and category set level, are available from Supplementary Table 1 and 2, respectively. For the precursor annotations, we curated family assignments, re-computed genomic clusters of miRNA genes, updated the chromosomal locations and source PubMed IDs for human, and added all similar categories for mouse and rat. All species are annotated with a new category containing high confidence precursors according to miRBase criteria. For human data, we transferred the disease annotations from HMDD to the new major release v3. We added associations from MNDR to allow disease comparisons against HMDD, and incorporated functional RNA interactions from NPInter. Lastly, novel categories such as the cellular localisation of miRNAs and regulatory interactions between miRNAs and transcription factors were incorporated from RNALocate and TransmiR, respectively. For the mature miRNAs, comparable changes apply as for the precursors in the cases of miRBase, MNDR, NPInter, and RNALocate-derived categorie sets. The gap between annotations of miRNA properties and their function is filled by categories on target genes taken from miRTarBase. To facilitate target-based enrichment of molecular pathways or biological function, we computed enrichments on target genes of miRNAs using Gene Ontology and KEGG. As an alternative for end-users, pre-computed significant enrichments of miRNAs associated with pathways provided by miRPathDB were made available for analysis. As the data from miRPathDB already involves a statistical pre-filtering, we implemented a new list of expert categories to highlight the underlying differences. Manually curated classifications from miRandola about known circular or extracellular miRNAs complete the final category dataset. Supposedly, the substantially enlarged number of categories might increase the average runtime of our algorithms, especially for the computationally intensive GSEA. Therefore, we profiled and improved our GeneTrail-based implementation to be three times faster, on average. [32].

Along with improving the data, we raised the available number of statistical parameter settings as well. First, users can request unadjusted or adjusted p-values using six published techniques to account for multiple hypothesis testing on the same dataset. In addition to the classical Bonferroni and Benjamini-Hochberg False discovery rate (BH-FDR) procedures, the adjustments proposed by Benjamini-Yekuteli, Hochberg, Holm, and Hommel can be selected. Moreover, the default behaviour of miEAA to correct p-values database / category set-wise was extended by a p-value pooling approach. In summary, the well-established alternatives for p-value correction can support highly customised research setups where alternate levels of stringency are required [33].

We also evaluated new visualisation features for the output of enrichment analyses to provide a simple overview and to improve comprehension. As a result, we made existing graphs interactive and implemented enrichment graphs with simulated background distributions for GSEA as well as automatic word cloud and heatmap plots for all enrichment algorithms. Word clouds display the names of obtained categories while scaling the size of the terms relatively to the number of hits that occurred and allow us to qualitatively compare the categories. On top of that, category to miRNA heatmaps depict log-transformed p-values at the combinations where hits occurred. This feature permits a simple way to compare the similarity of enriched / depleted categories with respect to associated miRNAs or precursors. The workflow of miEAA and example visualisations are displayed in Figure 1. Finally, we enhanced the general accessibility of miEAA through the implementation of a public API and a Python package, for which more details are provided below.

**Figure 1:**
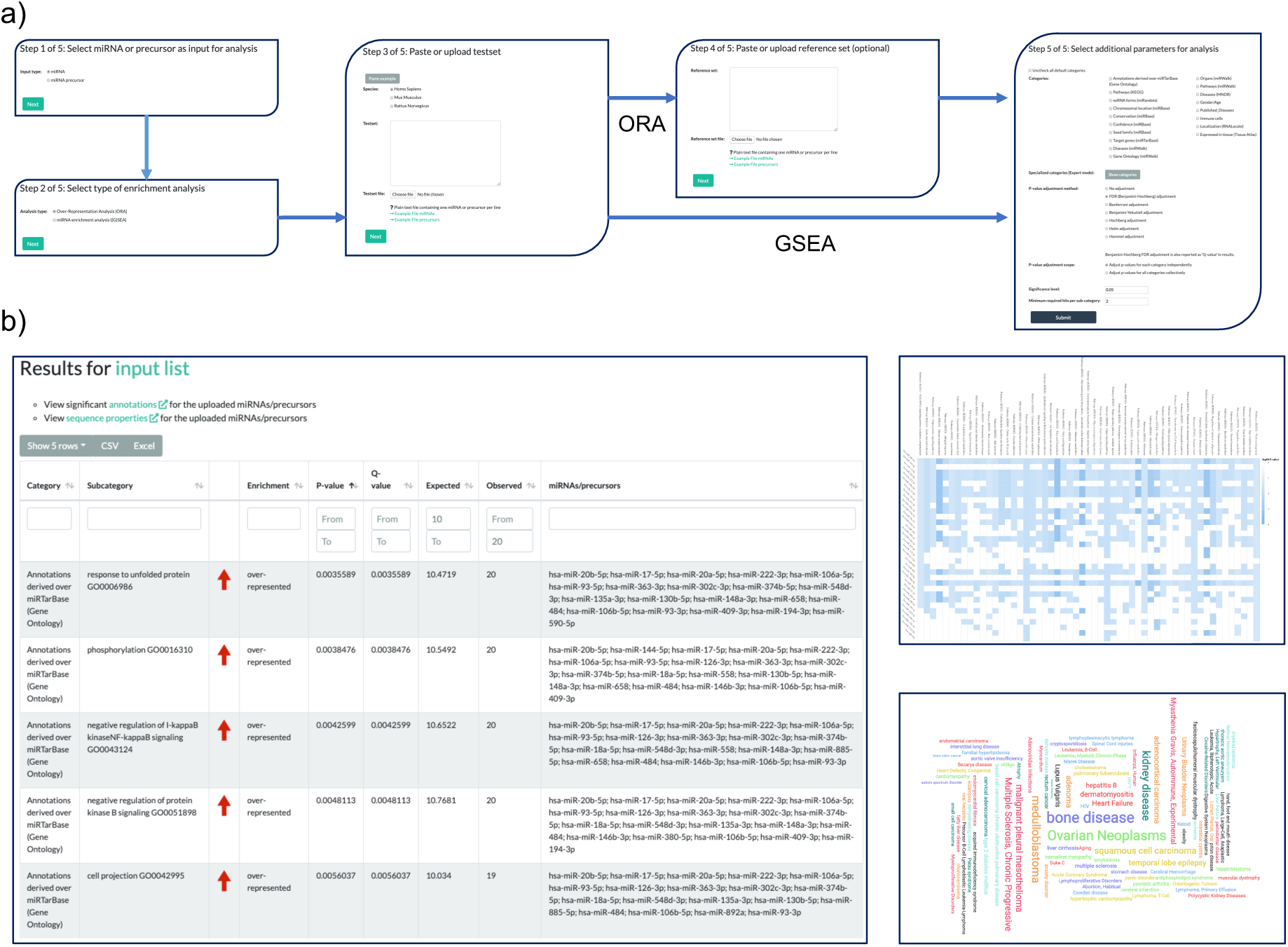
miEAA workflow and exemplary results. **(a)** Each miRNA / precursor enrichment analysis consists of at most five steps. First, users should select whether they want to perform enrichment on precursors or miRNAs. Second, the enrichment algorithm, i.e. either ORA or GSEA must be selected. Next, the desired test set can be defined either through a textbox or a file upload. The fourth step only appears for ORAs where custom background reference sets can be inserted or uploaded. This is optional since miEAA provides pre-computed reference sets for all categories. Lastly, the set of categories and databases as well as statistical parameters should be selected. **(b)** Typical result view for an ORA. Users can sort, select, filter, and export the obtained enrichment results interactively. Moreover, several visualisations of the results are provided for each run, such as the precursor / miRNA to category heatmap and the category wordcloud.

### Case study 1: Human kidney renal clear cell carcinoma

As the first case-study of miEAA 2.0, we acquired 591 human miRNA-seq samples from the kidney renal clear cell carcinoma (KIRC) project of TCGA, which can be divided into 520 Primary tumor (PT) and 71 Solid tissue normal (STN) samples. Sample information can be found in Supplementary Table 3. Of the 1, 881 precursors from miRBase v21, 321 are consistently detected in at least 50% of the samples for each biogroup. Among these, 282 were differentially expressed between PT and STN according to the FDR-adjusted wilcoxon test p-values (*p* < 0.01). Over-representation analysis of the precursors resulted in 541 significantly enriched and 7 significantly depleted (FDR-adjusted; *p* < 0.05) categories. As shown in Figure 2(b), a subset of miRNAs is ubiquitously present in significant categories, while others seem to be more specific. The top 10 categories sorted by increasing p-value are associated with cancer, including renal cell carcinoma. Also, the observed over expected ratio (123*/*48.6) indicates a strong enrichment (*p* = 2.80 × 10^−38^) of the de-regulated precursors with kidney and other types of cancer. A miRNA set enrichment analysis, using the list of detected precursors and sorted by effect size, revealed 253 enriched and 40 depleted categories. Here, the miRNA-precursor cluster 147, 189, 704 : 147, 284, 728 on the X chromosome is the most depleted category (*p* = 8.64 × 10^−10^), an observation that is in line with the depletion of precursor family hsa-mir-506. Interestingly, the list of highly enriched terms contains many transcription factors, the top 5 being *HEY1, WDR5, ELF1, BRD4*, and *FLI1*.

**Figure 2:**
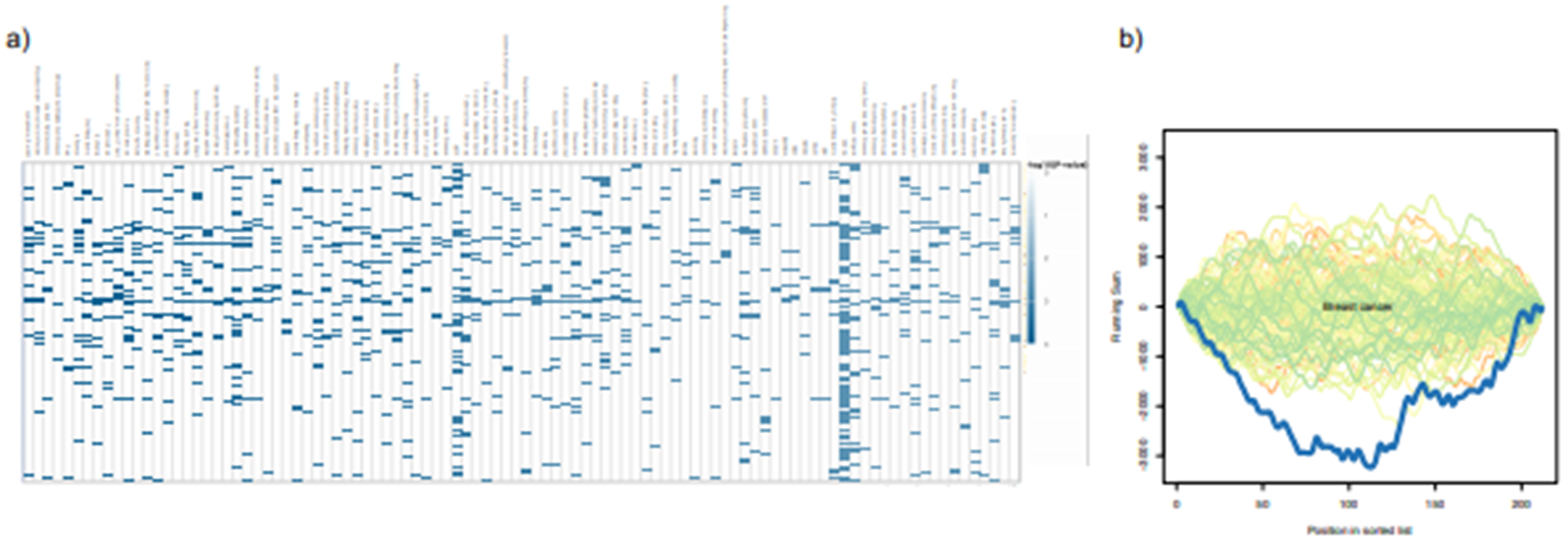
Web server visualisation of case study results. **(a)** Category to miRNA heatmap with −*log*10-scaled enrichment p-values from the first case study. **(b)** GSEA plot with simulated background distributions (green to orange lines) and actual depletion observed for breast cancer (dark blue line) during evaluation of the second case study.

### Case study 2: Mouse model for breast cancer progression

To showcase the novel support for model organisms in miEAA, we selected a dataset from GEO where circulating miRNAs from a breast-cancer mouse model were measured with microarrays [34]. The dataset comprises in total 36 samples from mutation-carrier (NeuT+) and age-matched wildtype (NeuT-) mice that were collected at the premalignant, preinvasive, and invasive stages of the disease. In this particular study, agilent microarrays probed with miRNAs from miRBase v19 were used on mice’s plasma extracted RNA samples. Sample information can be found in Supplementary Table 4. Following a detection threshold procedure similar to our first case study, 212 miRNAs remained for differential expression analysis. Of these, mmu-miR-6243 had to be discarded as a result of mapping the identifiers from miRBase v19 to v22.1, which we performed with the miEAA miRBase version converter. Subsequently, we applied GSEA on the list of miRNAs sorted by decreasing effect size between the premalignant and the invasive stage, for NeuT+ and NeuT-samples separately. Strikingly, the former run returned 311 significant categories, while the latter returned none. Overall, many more categories seemed to be depleted (*N* = 301) than enriched (*N* = 9), suggesting a wide-spread up-regulation of molecular pathways by miRNAs being down-regulated in NeuT+. For example, we found Macrophage differentiation (*p* = 2.54 × 10^−5^), Vasculature development (*p* = 1.60 × 10^−4^), and VEGF signaling pathway (*p* = 0.0016) to be depleted, which might be a signal for the increased tumor burden of NeuT+ mice at the invasive breast cancer stage. Moreover, we evaluated GSEAs for the comparison of NeuT+ and NeuT-at all three stages. While the first two setups returned a rather unspecific set of categories with all p-values located close to the significance boundary, the last comparison yielded many interesting results. First, observations were in line with the group-wise comparison along the age dimension, because all categories are depleted, i.e. no enrichments. Further, the results show that several dozen conserved miRNAs (*p* = 4.53 × 10^−5^) are down-regulated in the NeuT+ model at the invasive stage. More significant categories we found like exosome (*p* = 2.31 × 10^−5^) and circulating (*p* = 0.0086) miRNAs, breast cancer (*p* = 0.0094, Figure 2(b)), microRNAs in cancer (*p* = 0.028), and PI3K-Akt signaling pathway (*p* = 0.028) can be associated with this exemplifying study.

### New data export and browsable API

All data, results, and interactive plots shown on the web server are exportable to common data formats. Also, we were seeking to support the trend towards the development of reproducible and automated data analysis pipelines [12]. To this end, miEAA hosts a public, browsable API offering the same functionality as the web site, allowing one to access the miRNA converters and statistical algorithms remotely. This functionality is further augmented by a full-feature Python package with API library code and a command-line interface (CLI). For example, a regular workflow as performed on the website can be accomplished with three sequential calls to the web API or one call to the CLI. We provide code examples in the common data science programming languages Python and R to demonstrate this use-case. We also implemented the interface to solve two recurring problems in biological data analysis. First, reproducibility of statistical experiments can be improved, because usage of the versioned API in the context of a workflow manager such as Snakemake [23] or Nextflow fosters self-documenting research setups [35]. Second, oftentimes the analysis of miRNA high-throughput data involves the comparison of multiple biogroups, timepoints or other annotation variables. With the aid of our API and the package, multiple runs of miEAA can be performed at ease while minimising the time spent for set up and results aggregation.

## DISCUSSION

Statistical tools for biological enrichment analysis are a key to understanding data from high-throughput omics assays. However, the performance primarily depends on the quality of the underlying annotations and the statistical soundness. We show that new developments in the miRNA research field yielded an unprecedented set of biological categories, covering most aspects of miRNA properties and function, with cross-species analysis becoming increasingly important. On the other side, as with every statistical framework applied on biological data, assumptions are not always met and findings should be assessed critically in the light of further validation experiments. The novel release of miEAA attempts to cover these aspects by enhancing the set of available categories both quantitatively and qualitatively as well as through offering more (stringent) approaches for p-value correction. Also, a major limitation of some datasets concerns the availability of mature miRNA identifiers, as only precursor names were available from source databases. However, especially in the context of diseases, mature miRNA resolution is preferable to match the biological selectivity for one major miRNA arm being expressed. Datasets incorporated in miEAA were compiled either automatically or manually. TAM, another miRNA enrichment tool with functionality similar to miEAA, uses a fewer number of high-quality annotations, which come exclusively from manual curation [13]. A detailed comparison with respect to 22 criteria between our tool and TAM is shown in Supplementary Table 5.

We have demonstrated the capability of miEAA to yield novel biological results in cancer research. For the kidney renal clear cell carcinoma case study, we found a depletion of the mir-506 precursor family, which has been observed before in other types of cancers [36, 37]. Many interactions to transcription factors were also found for the up-regulated miRNAs, suggesting an increased regulatory burden due to the exceedingly transcriptional up-regulation observed in cancer. For example, HEY1, which is a transcriptional repressor has been characterised to be up-regulated in renal cell carcinomas [38]. For the mouse breast cancer progression study, we illustrated the backwards compatibility of miEAA with respect to miRBase. The overall observed depletion of pathways in mice agrees with our first case study. Moreover, the significant categories like vasculature development that are associated with morphogenesis, resemble an increased tumor burden of NeuT+ mice, which was previously confirmed with a large human RNA-seq dataset on breast cancer [39]. In both case studies we observed many associations with other types of cancers or diseases. While this may speak for a molecular and biological similarity, a certain publication bias, e.g. for cancer, is a confounding factor that skews the statistics [13].

Finally, we sought to improve accessibility of miEAA and develop a web-API in combination with a Python package and code examples. These features can also enhance its usability in other applications for miRNA research, for example to annotate functional sub-graphs in regulatory network analysis [40]. In conclusion, miEAA 2.0 is a flexible, comprehensive, and highly accessible tool for high-throughput miRNA annotation and enrichment analysis.

## DATA AVAILABILITY

miEAA 2.0 is freely available at https://www.ccb.uni-saarland.de/mieaa2. No login is required. Example code for API-usage and a pre-compiled Python package are freely available from https://github.com/Xethic/miEAA-API.

## SUPPLEMENTARY DATA

Supplementary Data are available online.

## ACKNOWLEDGEMENTS

We thank the authors of the utilised GEO dataset for providing their microarray samples to the general public. The results shown here are in whole or part based upon data generated by the TCGA Research Network: https://www.cancer.gov/tcga. We would like to express gratitude towards all specimen donors and research groups involved in the sample acquisition.

