## Supplementary Document for "miEAA 2.0: Integrating multi-species microRNA enrichment analysis and workflow management systems"

**Supplementary Table 1.** Databases and other resources used to generate miRNA/precursor categories for analysis in miEAA. Shown are the number of categories per species and resource.

**Supplementary Table 2.** Counts for category sets derived from the different data sources. A detailed breakdown of the category counts per input type, i.e. precursor or miRNA and species are displayed.

**Supplementary Table 3.** Sample metadata sheet from TCGA used for the kidney clear cell renal cell carcinoma case study.

**Supplementary Table 4.** Sample metadata sheet from GEO used for the breast cancer progression of a mouse model case study.

**Supplementary Table 5.** Detailed feature comparison between web servers miEAA 2.0 and TAM 2.0.
